# Rapid and Effective Inactivation of SARS-CoV-2 by a Cationic Conjugated Oligomer with Visible Light: Studies of Antiviral Activity in Solutions and on Supports

**DOI:** 10.1101/2021.10.18.464882

**Authors:** Kemal Kaya, Mohammed I. Khalil, Benjamin Fetrow, Hugh Fritz, Pradeepkumar Jagadesan, Virginie Bondu, Linnea Ista, Eva Y. Chi, Kirk S. Schanze, David G. Whitten, Alison M. Kell

## Abstract

This paper presents results of a study of a new cationic oligomer that contains end groups and a chromophore affording inactivation of SARS-Cov-2 by visible light irradiation in solution or as a solid coating on wipes paper and glass fiber filtration substrates. A key finding of this study is that the cationic oligomer with a central thiophene ring and imidazolium charged groups give outstanding performance in both killing of *E. coli* bacterial cells and inactivation of the virus at very short times. Our introduction of cationic N-Methyl Imidazolium groups enhances the light-activation process for both *E. coli* and SARS-Cov-2 but dampens the dark killing of the bacteria and eliminates the dark inactivation of the virus. For the studies with this oligomer in solution at concentration of 1 μg/mL and *E. coli* we obtain 3 log killing of the bacteria with 10 min irradiation with LuzChem cool white lights (mimicking indoor illumination). With the oligomer in solution at a concentration of 10 μg/mL, we observe 4 logs inactivation (99.99 %) in 5 minutes of irradiation and total inactivation after 10 min. The oligomer is quite active against *E. coli* on oligomer-coated wipes papers and glass fiber filter supports. The SARS-Cov-2 is also inactivated by the oligomer coated glass fiber filter papers. This study indicates that these oligomer-coated materials may be very useful as wipes and filtration materials.

## INTRODUCTION

The World Health Organization (WHO) has declared the novel coronavirus outbreak of 2019 to be a global pandemic. To date, severe acute respiratory syndrome coronavirus 2 (SARS-CoV-2) has infected more than 229 million people globally, and claimed more than 4 million deaths in the world. ^1^ Several vaccines have been developed that are very effective against SARS-CoV-2. Although more than six billion doses of the vaccines have been administered, a high percentage of the world population has been reluctant to take the vaccines, despite scientific evidence of their effectiveness. It appears that getting a sufficient fraction to be vaccinated and thus obtain “herd” immunity is difficult and likely impossible for the current SARS-CoV-2 and future viral pathogens.

The virus is known to spread via respiratory droplets^2^, close contact with infected individuals, and by viral contamination of frequently touched surfaces. ^3^ As such, government and health authorities have urged the use of personal protective equipment (PPE), such as masks, the avoidance of indoor congregation, and disinfection of surfaces.^4^ While mask wearing has been shown to mitigate the spread of Covid-19, there is also much resistance to the wearing of masks and closure of schools and popular venues such as theaters, restaurants and businesses. There is a clear need for antimicrobial agents that inactivate SARS-CoV-2 and minimize its spread via surface contact and air. Several recent works have shown new methods for disinfecting surfaces against coronavirus infections.^5^ In a study published in 2020, Poon, Ducker, et al. reported Cu_2_O bound with polyurethane rapidly reduce virus titer and prevent re-infection for periods of days to weeks.^6^ Other studies indicate the virus can be inactivated but in many cases the components are volatile and cannot provide lasting disinfection.^4^ Currently, there are few treatment options and very few long-lasting disinfectants available for inactivating the virus before it can spread and infect humans. While masks and protective clothing and “social distancing” may offer some protection, their use has not always halted or slowed the spread.

Our group has studied oligomers of poly-phenylene ethynylene ^7, 8, 9^ as potent antimicrobial agents^10,11,12^ against bacteria, fungi and viruses.^13,14,15,16^ Specifically, we have found that the non-enveloped viruses attacking *E. coli* exhibit potent antiviral activity under both light-activation and dark treatment with several cationic oligomers and polyelectrolytes.^14^ For both dark and light-activation the level of inactivation reaches several logs.^14^ We have recently developed a series of antimicrobial reagents that are effective against SARS-CoV-2 in aqueous suspensions when activated by ultraviolet or visible light.^16^ These materials include a series of cationic poly-phenylene-ethynylenes and polythiophenes and smaller cationic and anionic oligomers.^16^ In contrast to our earlier findings these materials only exhibited virus inactivation under visible or uv light irradiation.^16^ We attributed the lack of dark antiviral application to the viral envelope of the SARS-Cov-2. While it appears that the dark activity involving membrane penetration or destruction cannot occur with these materials, they must engage in a non-damaging ground state docking process that enables singlet oxygen formation from the excited triplet of the oligomer or polymer and subsequent penetration of singlet oxygen into the virus core. Subsequent to publishing our first paper we have focused our attention to developing stronger oligomeric antimicrobials, particularly on their speed of inactivation^17,18,19^ and employment in other formats.^20,21,22^ Herein, we report the development of a new oligomer (**1**) that we have found to very rapidly and effectively inactivate SARS-CoV-2 in aqueous solutions with visible light where these materials are soluble and in combination with scaffolds in filtration trials which offer the potential for water and air disinfection processes. In this paper we discuss a method of destroying, inactivating, reducing the infectivity of, or otherwise inhibiting the activity of SARS-CoV-2 by irradiation of **1** with visible light.

## RESULTS and DISCUSSION

### Rationale and synthesis of new OPE compounds

In follow up to our initial studies, we tested new oligomers and polymers with SARS-CoV-2 and found that two of five materials tested with visible light activation (“cool white” light tubes from LuzChem) were very effective in inactivating the virus with these lamps, which are described as a good model for interior lighting by artificial light sources. ^16^ In designing new oligomers for potential applications against SARS-CoV-2 we considered that the simplest oligomers we found effective were an end-only OPE (EO-OPE-1) where the chromophore and charged groups may be easily tuned by straight-forward synthetic methods. We designed and synthesized two OPEs substituted with a central thiophene ring and potent but different charged end groups. These structures are shown in **Scheme** 1 below.

**Scheme 1:**
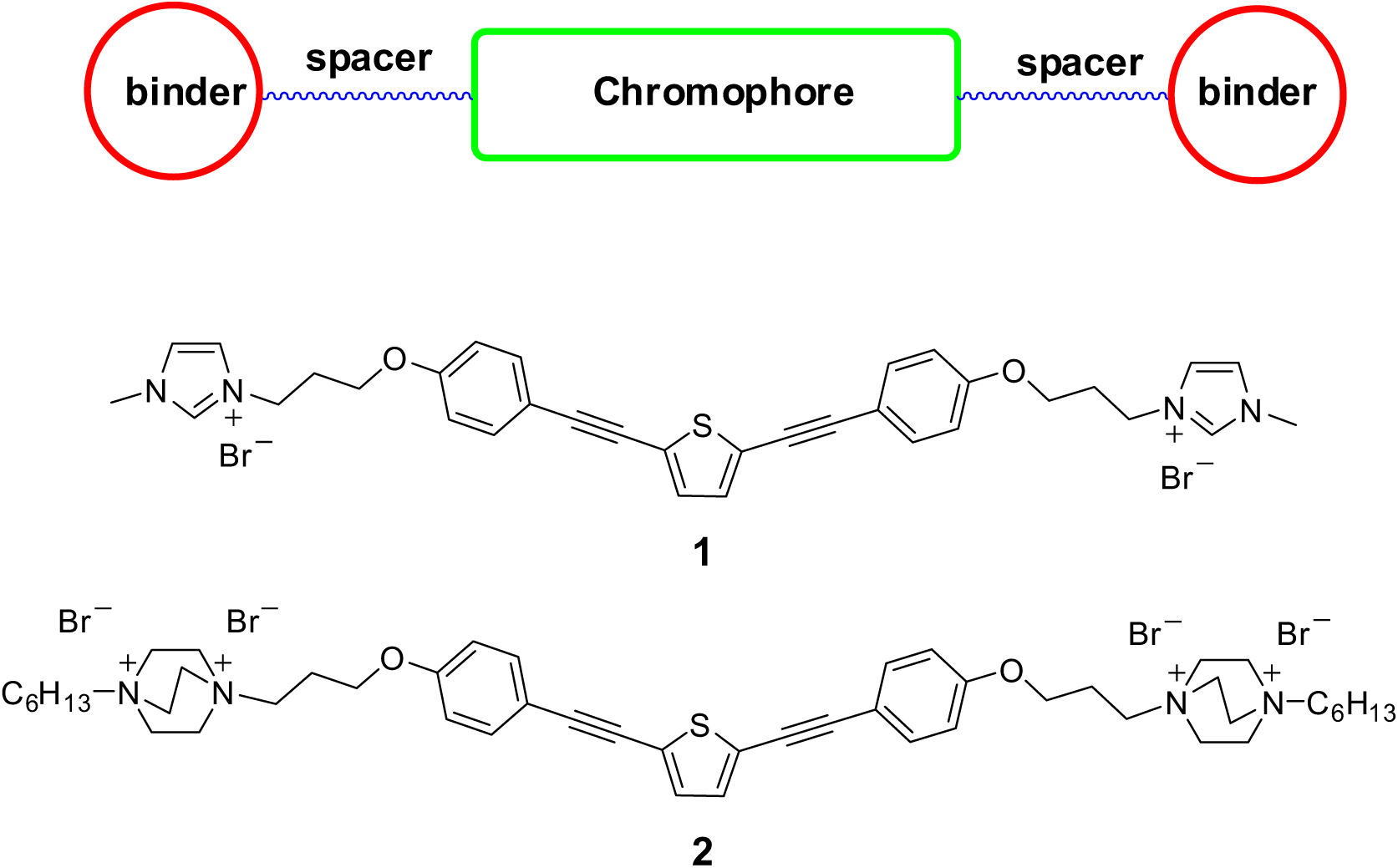
Top: functional components of end-only functionalized oligomers. Structures of **1** and **2** used in this study are shown below.

While we could synthesize each of these oligomers from a common dibromo derivative, we found that the DABCO derivative **2** was impractical for use due to its insolubility in water. However, oligomer **1** is readily soluble in water and easy to use. Before testing **1** with SARS CoV-2 we decided to test **1** with *E. coli* in different formats including solutions of **1**, supported samples of **1** adsorbed on different solids including wipes materials and other potential scaffolds.

### Antibacterial activities of compound 1 against *E. coli* in solution

Gram-negative *E*.*coli* were investigated under cool white light irradiation at 1 μg/mL and 10 μg/mL concentrations in order to minimize potential inner filter effect^23,24^ and for different times (5, 10 and 30 min). Log reduction or “log kill” was determined by serial dilution assay where colony forming units were quantified. The results are summarized in Figure 1 and Table 1. Noticeably, for all control experiments, there is no killing activity in dark (control dark) and no killing activity from the light that is used (control light).

**Table 1.** Summary of antimicrobial activity of compound **1** in solution and on supporting materials.

| Concentration | Format | Log Kill | Irradiation Time |
| --- | --- | --- | --- |
| 10 $\mu$ g/mL | Solution | 4 log | 5 min |
| 10 $\mu$ g/mL | Solution | 6 log | 10 min |
| 1 $\mu$ g/mL | Solution | No Killing | 5 min |
| 1 $\mu$ g/mL | Solution | 3 log | 10 min |
| 1 $\mu$ g/mL | Solution | 6 log | 30 min |
| 0.1 $\mu$ g/mL | Solution | No Killing | 30 min |
| 10 $\mu$ g/mL | Wipe Paper | 5 log | 30 min |
| 10 $\mu$ g/mL | Glass Fiber Filter<br>#2 | 5 Log | 30 min |

**Figure 1.**
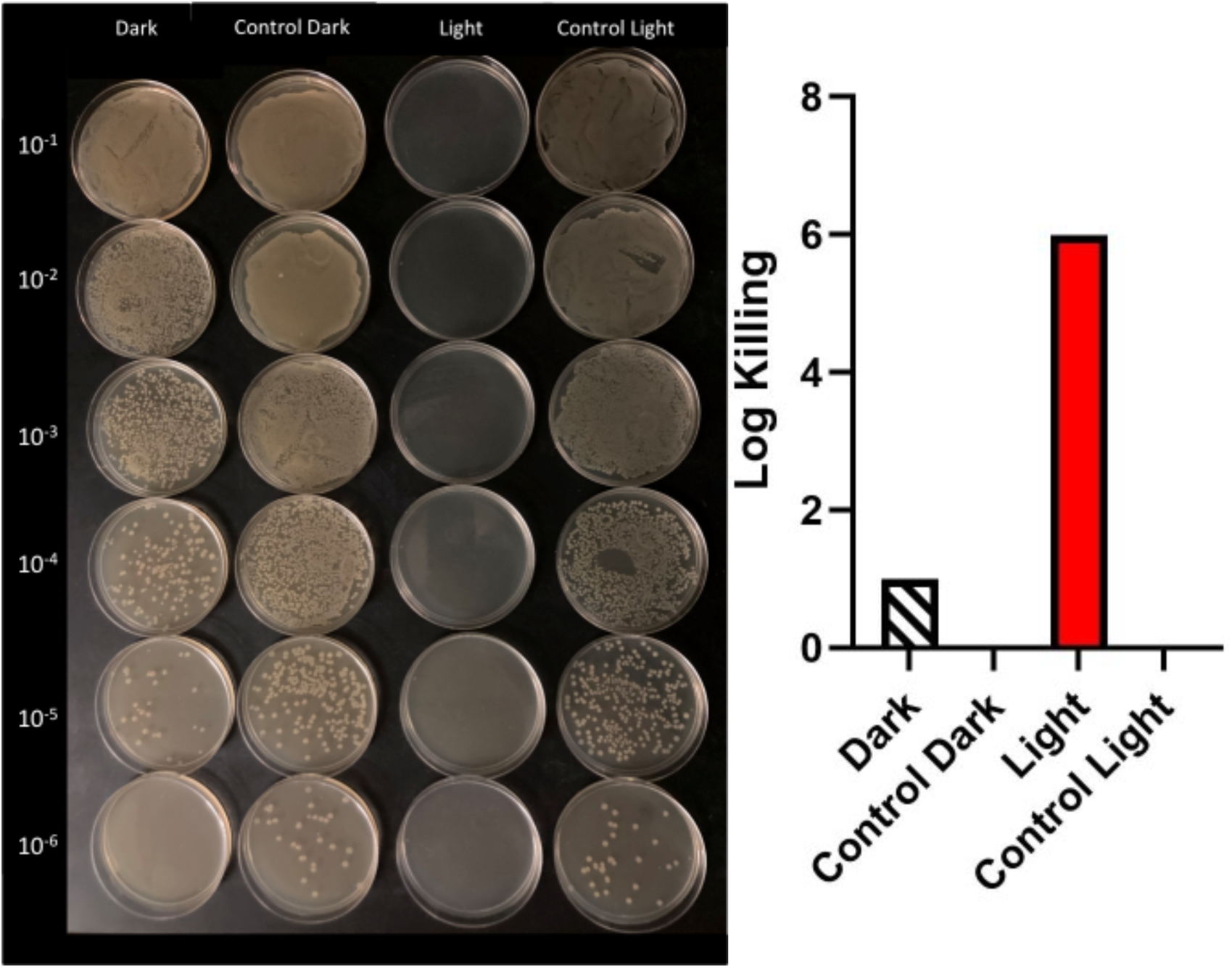
Cell counting of serial dilutions of *E. coli*. Under irradiation of 10 μg/mL of compound **1** with visible light and in dark (above, left) and graphical representations of log kills in dark and light. There was no killing on irradiation (Control Light) and dark control (Control Dark) samples.

**Figure 2.**
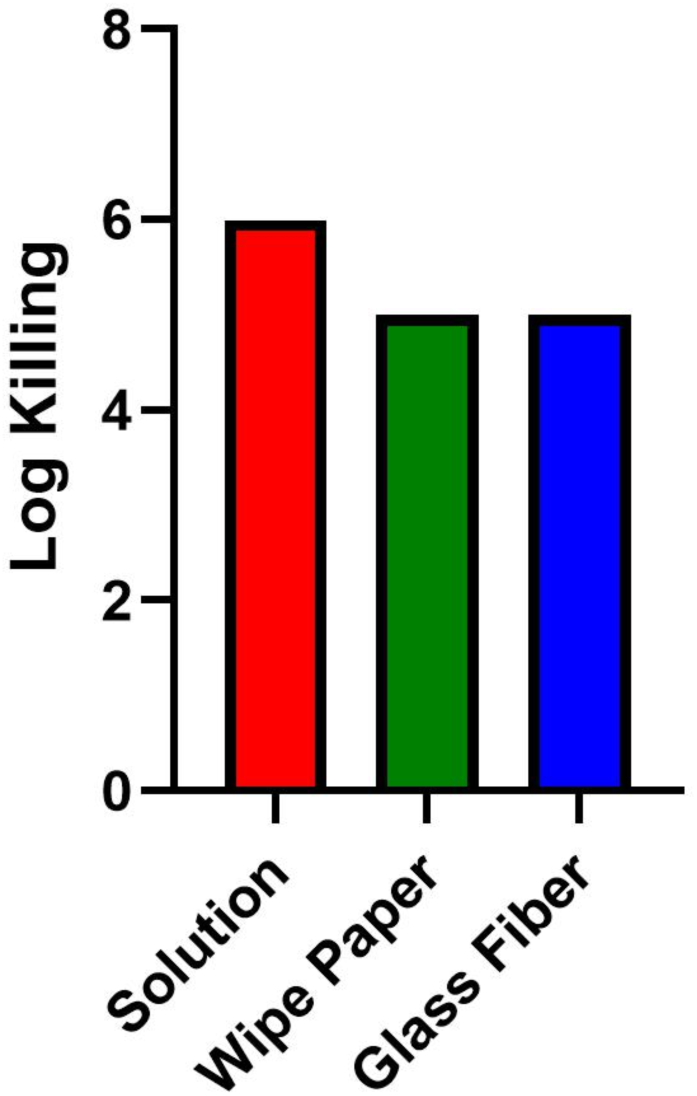
Antimicrobial activity of compound **1** against *E. coli* upon exposure to cool white light for 30 min.

When 10 μg/mL concentration of the compound was used, there is very little killing under dark conditions. However, compound **1** is extremely reactive against *E. coli* and exhibits enhanced antimicrobial activity when it is irradiated with cool white light. We have observed 4 log kills after 5 min irradiation (SI Figure S1) and 6 log kills for 10 min irradiation Figure 1. To the best of our knowledge, **1** is the only compound of the OPE family that shows 5 log killing enhancement with light irradiation. Interestingly, when we have run the biocidal experiment at 1 μg/mL concentration for 5, 10 and 30 min, dark reactivity of compound **1** has disappeared.

### Antibacterial activities of compound 1 against *E. coli* on supporting wipe and glass fiber materials

Having demonstrated effective antimicrobial activity of **1** in solution, we decided to coat commercial wipes paper and glass fiber filters with **1** and test them for biocidal activity. These materials were coated with **1** and the coated materials were cut so that the final effective concentration of 1-10 μg/mL was used for biocidal experiments. A piece of glass fiber (3 × 6 cm^2^, 127 mg) was cut and immersed into 50 mL of conjugated cationic oligomer solution (100 μg/mL) for 12 hrs. After soaking, the filter was removed from solution and dried at room temperature for 3 days. UV Vis spectra was used to determine how much compound was absorbed by glass fiber. Because of the high absorbance of compound **1** at 100 μg/mL, this solution was diluted to 10 μg/mL and UV spectra were taken (SI Figure S2). Considering the dilution, calculations were done accordingly in order to determine the amount of compound **1** that is absorbed by glass fiber. To test whether or not coated glass fiber is releasing **1** into water or PBS, coated glass fiber was immersed into milliQ water and PBS. There was no detectible release by UV into either water or PBS.

The antibacterial activity of both coated samples shows 5 logs of inactivation, 1 log less inactivation than in solutions of **1**. When the materials were coated, it was found that there was no release of **1** by rinsing of the materials with water or PBS. Thus, we can conclude that **1** is effective against bacteria while adsorbed to the support and that the inactivation is not due to released **1** in the solution.

### Antiviral activity of compound 1 against SARS-CoV-2

For the studies of **1** vs SARS-CoV-2, we used a procedure similar to that described above. However only samples of **1** in solution added to aqueous suspensions of SARS-CoV-2 and samples of **1** coated on glass fiber filter mater were studied. Results (Figure 3) show that with compound **1** in solution mixed with a suspension of SARS-CoV-2, there is rapid and complete inactivation of the virus over a period of several minutes under cool white light illumination. For the one measurement (three determinations) where there was measurable viral activity (5 minutes) there was already ∼3 logs or 99.9% inactivation of the virus! This indicates that if compound **1** is deployed as a wipe where some of the oligomer is released into aqueous solution, there will be complete and rapid inactivation. The ideal wipe could inactivate all virus on a surface and deposit sufficient oligomer to inactivate any viruses contacting the surface for a prolonged time scale. The same would be true for a spray containing compound **1**.

**Figure 3.**
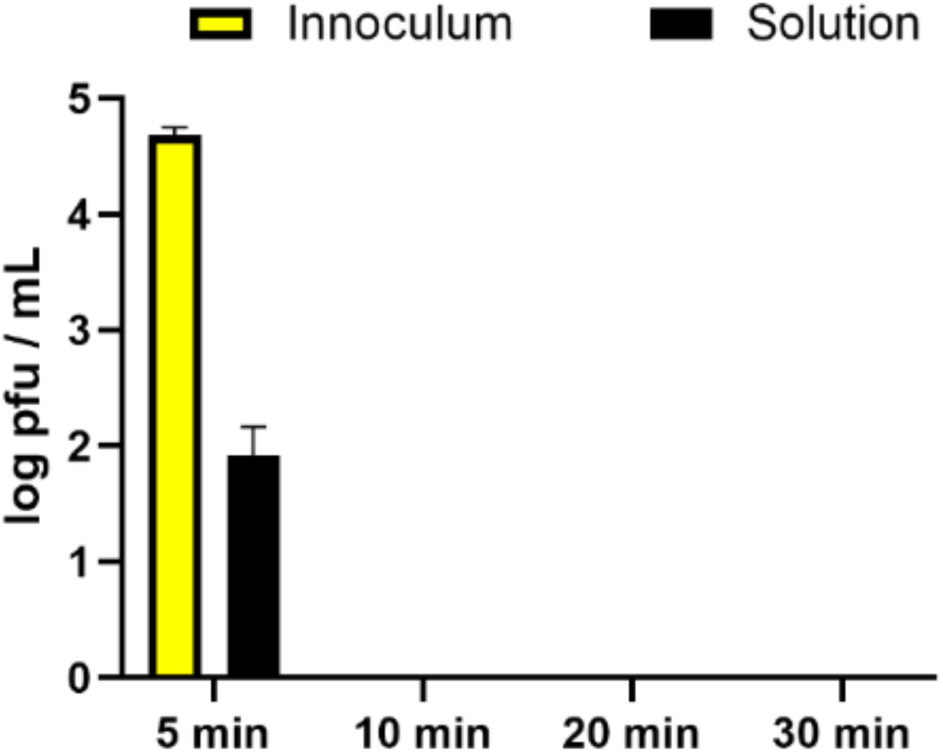
Antiviral activity of 10 μg/mL of compound **1** in solution against SARS-CoV-2. Compound **1** was incubated with 1× 10^5^ pfu/mL SARS-CoV-2 in “cool white” light for the indicated times. Viral titer was quantified by plaque assay on Vero E6 cells. Values shown are the average titer from at least three independent experiments (±SEM). Inoculum is a control sample containing untreated virus and was not irradiated. There is no active virus left after 10, 20 and 30 min irradiation.

The very rapid inactivation of the virus under visible light irradiation similar to indoor illumination suggests **1** should be very useful and effective in applications such as sprays or wipes against viruses on surfaces or as aerosols, even under ambient irradiation. We also tested **1** as a coating on glass fiber filtration material. Figure 5 shows both coated and uncoated samples of a glass fiber filter material provided by HYDAC (details). While the coating process takes significantly longer for this filter material than for conventional wipes materials (12 vs. 2 hours), the photographs in Figure 5 clearly show strong and uniform fluorescence from compound **1**. When the coated filter is immersed in a suspension of SARS-CoV-2 and irradiated with “cool white” light, there is a slower but complete inactivation of the virus. The results shown in Figure 6 indicate that incubating compound **1** with a sample filtration material (sample number 2 glass fiber 3.5 mfp)results in a filter material with a coating that is not removed by treatment with water or buffer solution, but is active against SARS-CoV-2 and results in ∼5 log reduction after 30 min irradiation with visible light in a stirred^1^ (but not under filtration) suspension. This result indicates that the combination of the HYDAC filter materials impregnated with compound **1** may provide a highly effective and efficient filtration system in which the virus is entrapped and destroyed.

The results presented above indicate that compound **1** is a powerful antiviral reagent that can be used in solution, sprays and wipes as an effective disinfectant against surfaces contaminated with SARS-CoV-2, and very likely against other enveloped viruses. The results shown in Figures 4 and 5 also suggest that residual **1** introduced on the same surfaces will provide active disinfection against new viruses. The results shown in Figure 5 suggest that **1**, when deposited on a filter material can provide effective decontamination of air or water samples in a filtration device.

**Figure 4.**
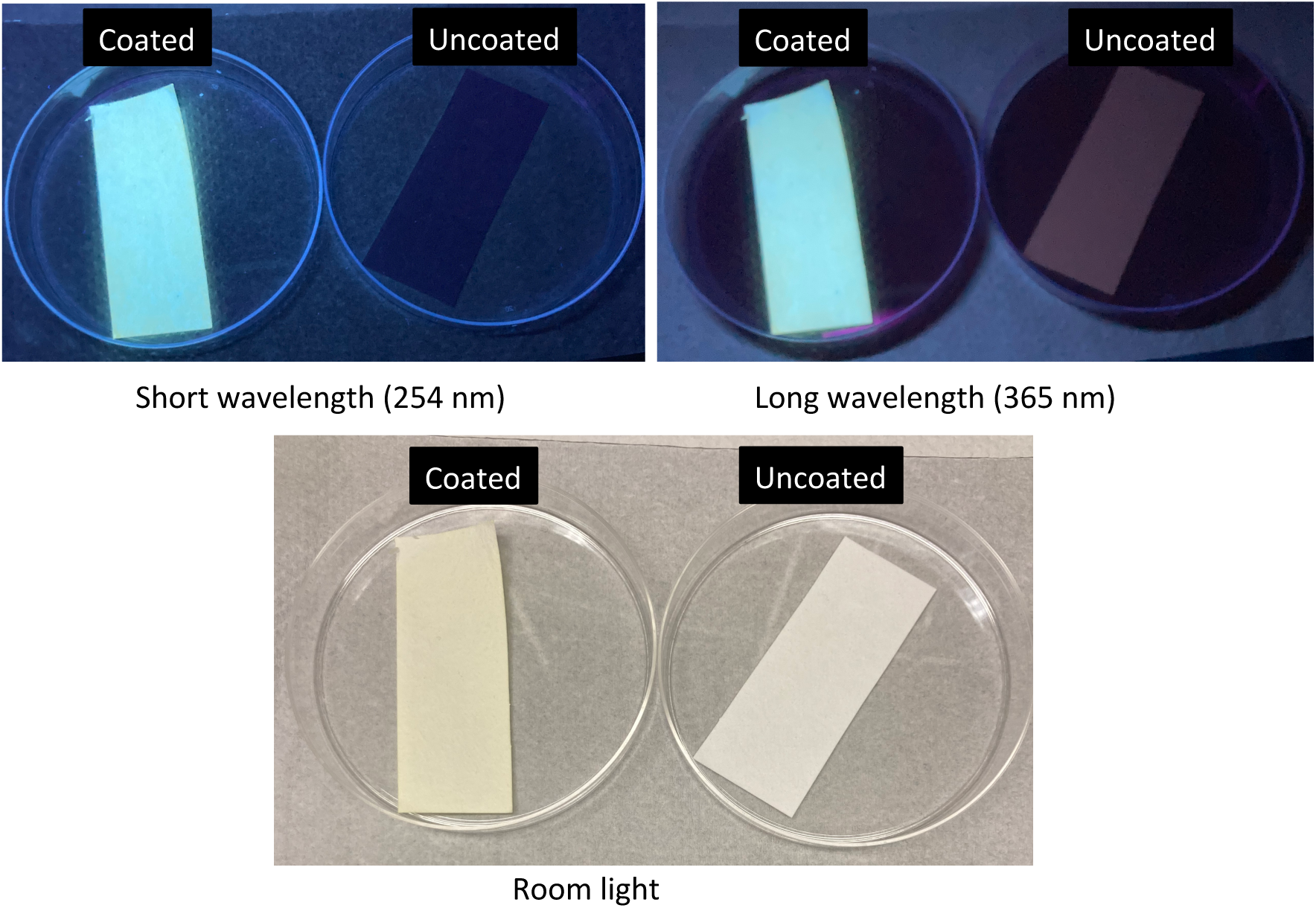
Photographs of uncoated and compound **1** coated glass fibers filters under different illuminations. In room light there is little difference between the filter samples. Under irradiation with both UV lamps, there is a striking difference in their appearance that we can attribute to fluorescence from compound **1**.

**Figure 5.**
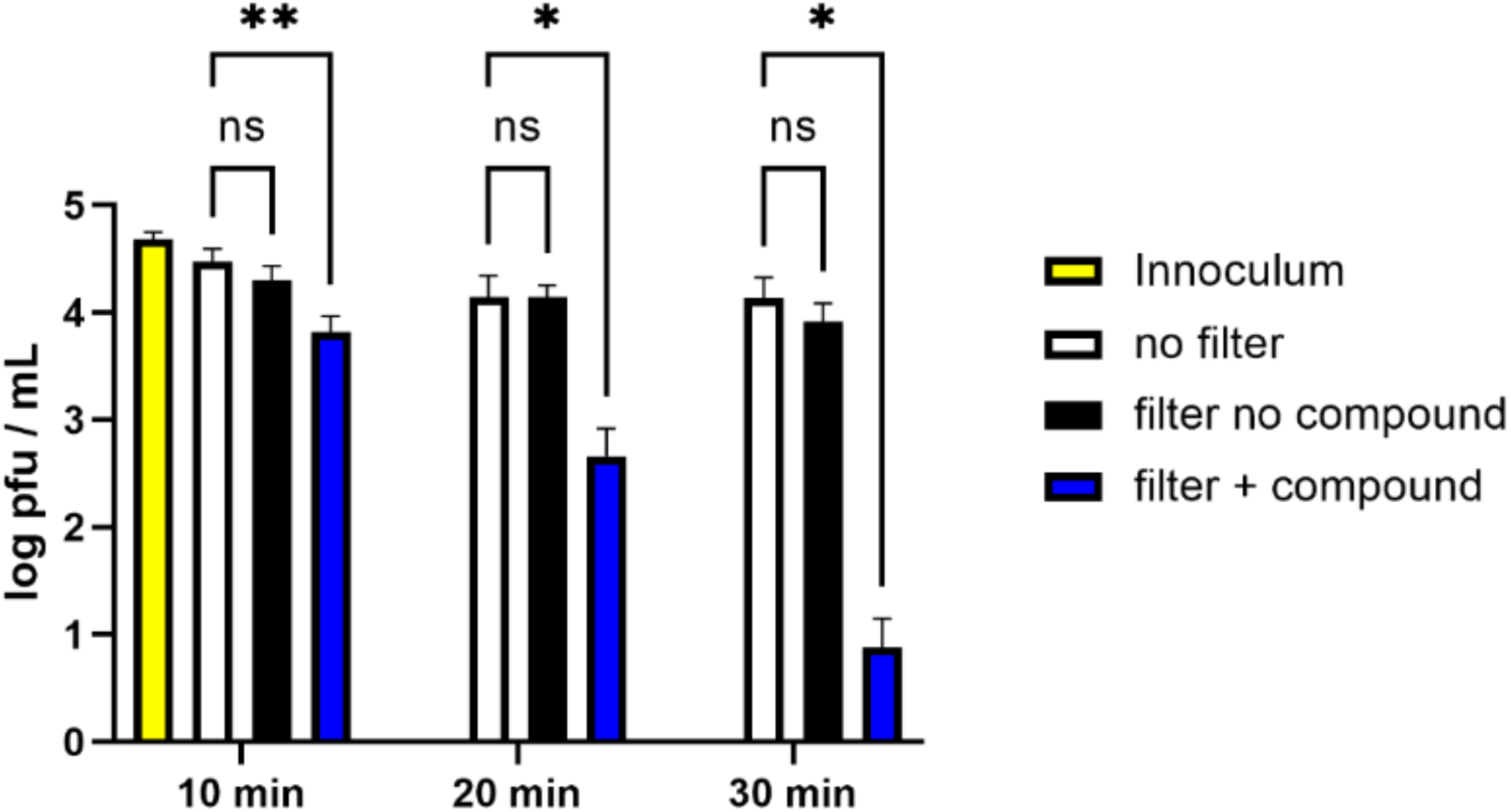
Antiviral activity of glass fiber filter coated with compound **1** against SARS-CoV-2. 1× 10^5^ pfu/mL SARS-CoV-2 was incubated with filter coated with compound **1** (filter + compound), uncoated filter material (filter no compound), or no filter in the presence of “cool white” light for the indicated times. Inoculum is a control sample containing untreated virus and sample was not irradiated. The amount of compound **1** present in the “filter + compound” samples is equivalent to 10 μg/mL in the sample solution. Viral titer was quantified by plaque assay on Vero E6 cells. Values shown are the average titer from at least three independent experiments (±SEM). * denotes p < 0.05, two-way ANOVA with multiple comparisons correction.

### Animal toxicity testing of compound 1 by Third Party Laboratories

We have recently supplied samples of compound **1** in solution to laboratories testing against toxicity against animal models. Preliminary results of the testing indicate that **1** is non-toxic in tests of eye irritation on rabbits, oral toxicity in rats, and nonirritating in dermal testing in rabbits. These tests are consistent with what we have found in tests conducted in our laboratories with several other phenylene ethynylene polymers and oligomers.^25,26,27^

## CONCLUSIONS

The results of the present study indicate that a new compound, an “End-Only” pligomeric phenylene ethynylene containing a central thiophene and cationic imidazolium charged groups (**1**) shows remarkable antiviral activity with visible light, inactivating SARS-CoV-2 with nearly three logs in five minutes in solution and up to almost 5 logs in a supported format on a glass fiber filtration material. These results provide strong evidence that materials incorporating this oligomer can be effective and practical for elimination of SARS-CoV-2 with visible light irradiation in solution or with wipes as filters and/or filtration devices. These materials and their deployment could help end the current pandemic and prevent future pandemics by intercepting the virus before it can propagate and spread.

## MATERIALS AND METHODS

### Synthesis of the compounds

Incorporation of trimethylsilyl acetylene groups at 2- and 5-positions was accomplished by Sonogashira coupling reaction in THF yielding 2,5-bis[ethynyl(trimethylsilyl)] thiophene (**3**). TMS groups were desilylated using K_2_CO_3_ in DCM/MeOH affording the desired 2,5-Bis(ethynyl)thiophene (**4**). Replacement of one Br atom on 1,3-dibromo propane with 4-iodo phenol yielded compound **5**. Coupling of compounds **5** and **4** by sonogashira reaction afforded compound **6**.^**Error! Bookmark not defined**., 28, 29, 30^ Replacements of Br atoms of compound **6** with N-Methyl imidazole gave compound **1** and with DABCO yielded compound **2**.

Synthesis of the compound is summarized in Scheme 2.

**Scheme 2.**
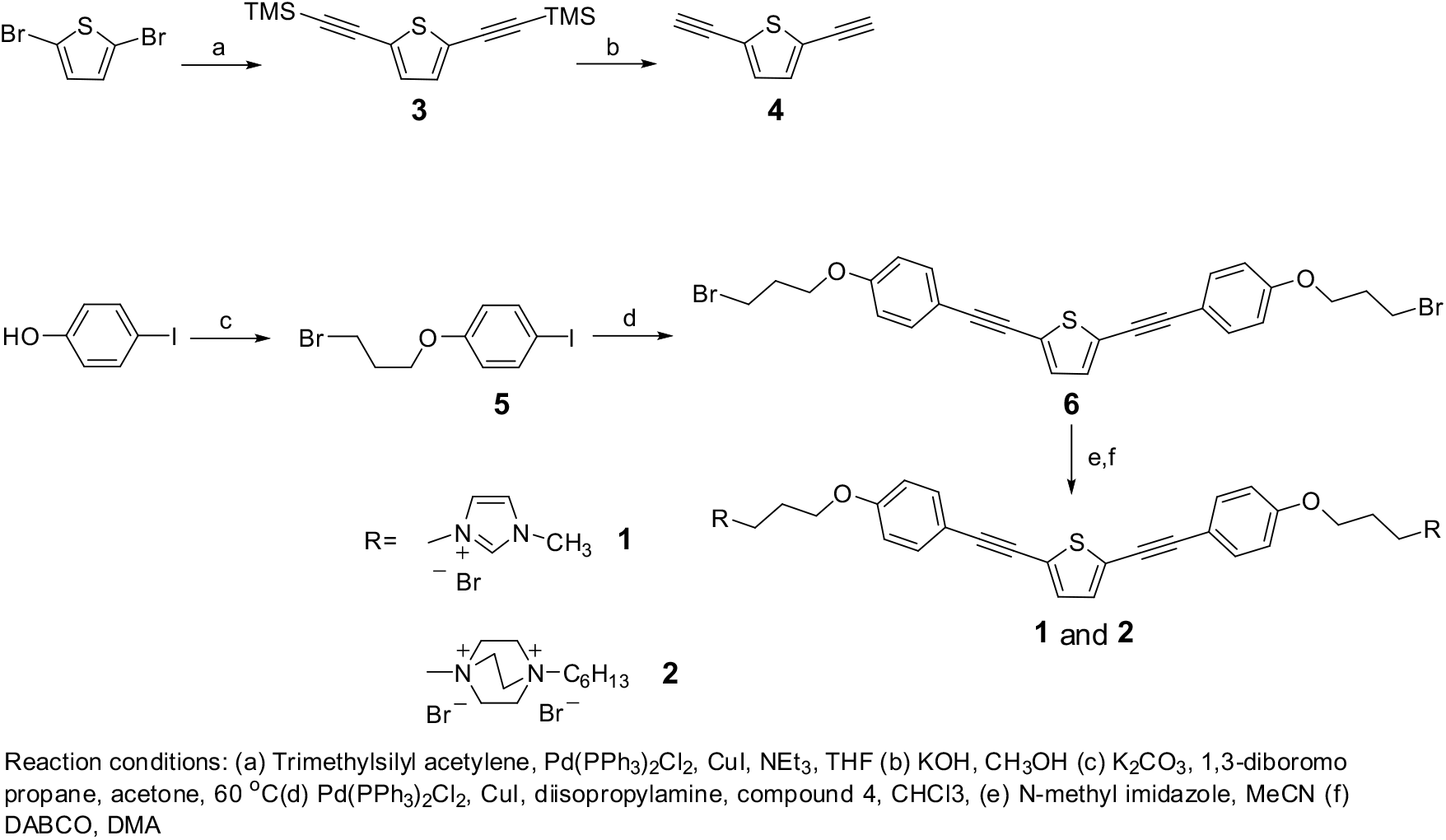
Synthesis of compounds 1 and 2.

### Preparation of oligomer coated glass fiber filters

A stock solution of compound **1** (100 μg/mL) was prepared by dissolving compound **1** in water. A piece of glass fiber filter (3 × 6 cm^2^, 127 mg, HYDAC company)) was cut and immersed into 50 mL of the OPE solution for 12 hrs. After soaking, the filter was removed from solution and dried at room temperature for 3 days. In order to calculate how much compound is absorbed by the glass fiber filter, UV-Vis spectra of the solution were taken before and after immersing and removing the filter (SI Figure S2). 0.84 mg of coated glass fiber filter was cut to use for a final oligomer concentration of 10 μg/mL in the SARS-CoV-2 plaque assay as described below to determine antiviral activity.

### Preparation of compound 1 coated commercial wipe paper

A stock solution of compound **1** (25 μg/mL) was prepared by dissolving the compound in water. A commercial wipe paper sheet (3 × 6 cm, Spontech) was cut and immersed into 30 mL of oligomer solution for 2 hr. After 2 hr, the paper was removed from solution and dried at room temperature for 3 days. In order to calculate how much compound is absorbed by the wipe paper. UV-Vis spectra of the solution were taken before and after immersing paper (SI Figure S3). 1.6 mg of coated wipe (2 × 2 mm) was cut for a concentration of 10 μg/mL for subsequent testing.

### Bacteriocidal experiments

All bacterial studies were carried out under sterile conditions (flaming, alcohol rinse, etc.). *E. coli* cells (ATCC 29425) were obtained from ATCC and used to generate a stock culture stored in 20% glycerol at -70 ° C. *E. coli* cell colonies were grown in Luria-Bertani (LB) broth. Biocidal testing entailed the scraping colonies off their agar plates and transferring them to nutrient broth for growth. Cells were then incubated in an orbital incubator shaker (American Instruments, Lafayette, CA) for approximately three hours at 37 ° C to the exponential growth phase (O.D. ∼0.6 at 600 nm). *E. coli* cells were then collected by centrifugation and washed three times with phosphate buffer saline (PBS). Supernatant was removed and discarded. The remaining pellet was resuspended in 7.2 mL of PBS solution.

Six one-dram vials were labeled as Light, Dark, Control Light, Control Dark, Cell Control Light. Cell Control Dark. 1.6 mg of coated wipe paper pieces were put into the Light and Dark labeled vials. For control experiments, Control Light and Control Dark labeled vials had 1.6 mg uncoated wipe papers. Cell Control Light and Cell Control Dark vials did not have any coated or uncoated wipe papers. To each vial, 900 μL of cell suspension and 100 μL of sterile PBS were added (final concentration of 10 μg/mL of the coated oligomer). All light labeled vials were irradiated for 30 minutes with 10 LZC-Vis lamps in a photoreactor (LZC-ORG, Luzchem Research Inc., Ottawa, Canada) whereas respective dark labeled vials were stored for the same time in a light protected box at room temperature. After irradiation, the experiment was stopped by diluting the cells 10 times with PBS. Further dilution steps up to 1 × 10^6^ were performed, and 100 μL of each dilution step was spread on an agar plate. After 18 hr of incubation at 37 °C, the plates were examined and the number of colonies was counted for each sample.

### Mammalian cells and SARS-CoV-2 viruses

Vero E6 cells (ATCC, CRL-1586) were cultured in Dulbecco’s modified Eagle’s medium (DMEM) supplemented with 10% heat-inactivated FBS, 1% pen/strep, 2 mM L-glut, 1% non-essential amino acids, and 1% HEPES. Severe acute respiratory syndrome coronavirus 2 (SARS-CoV-2, Isolate USA-WA1/2020) was acquired from BEI Resources (NR-52281) and propagated in Vero E6 cells for 3 days. Infectious virus was isolated by harvesting cellular supernatant and spinning at 1000 rpm for 10 minutes to remove cellular debris and stored at -80°C. SARS-CoV-2 infectious particles were quantified in media by standard plaque assay. Briefly, Vero E6 cells were treated with serial dilutions of SARS-CoV-2 in DMEM (4% FBS) and incubated at 37°C for 1 hr. An overlay of 2.4% cellulose (colloidal microcrystalline, Sigma #435244) and 2x DMEM, 5% FBS was added to each well and incubated for 3 days at 37°C, 5% CO_2_. On day 3, cells were fixed with 4% paraformaldehyde and stained with crystal violet to visualize and count the number of plaque forming units (pfu).

### Inactivation of SARS-CoV-2 by compound 1 in solution

A solution of compound was diluted to 10 μg/mL in DMEM culture media (4% FBS). SARS-CoV-2 was added to compound-media solution at a final concentration of 1× 10^5^ pfu/mL. Virus was incubated in media supplemented with a volume of RNA/DNAse-free water equal to that used for the compound dilutions served as a negative control. Samples were then incubated at room temperature inside a LuzChem light chamber and exposed to the “cool white” light for the specified time intervals. Infectious virus present in the samples was then quantified by plaque assay.^16^

### Inactivation of SARS-CoV-2 by compound 1 coated glass fiber

For experiments testing compound-coated filter inactivation, a media solution of SARS-CoV-2 at a concentration of 1× 10^5^ pfu/mL was added to 1.5 mL tubes containing **1** coated filter at a solution concentration of 10 μg/mL and gently agitated by pipetting up and down multiple times. As a negative control, filters not coated with **1** were incubated with viral preparations and treated similarly. Samples were then incubated at room temperature inside a LuzChem light chamber and exposed to “cool white” light for the specified time intervals. Infectious virus present in the samples was then quantified by plaque assay.

## Notes

The authors declare the following competing financial interest(s): KSS and DGW have interest in a start up company, BioSafe, LLC that has licensed patents related tothe materials that are the focus of this work.

## ACKNOWLEDGMENTS

A.M.K. was supported by NIH (grant 1K22AI141680-01A1). K.S.S. acknowledges the Welch Foundation for support (grant no. AX-0045-20110629). The authors thank HYDAC Corporation and Spuntech Industries Inc. for samples.

## Graphical Abstract

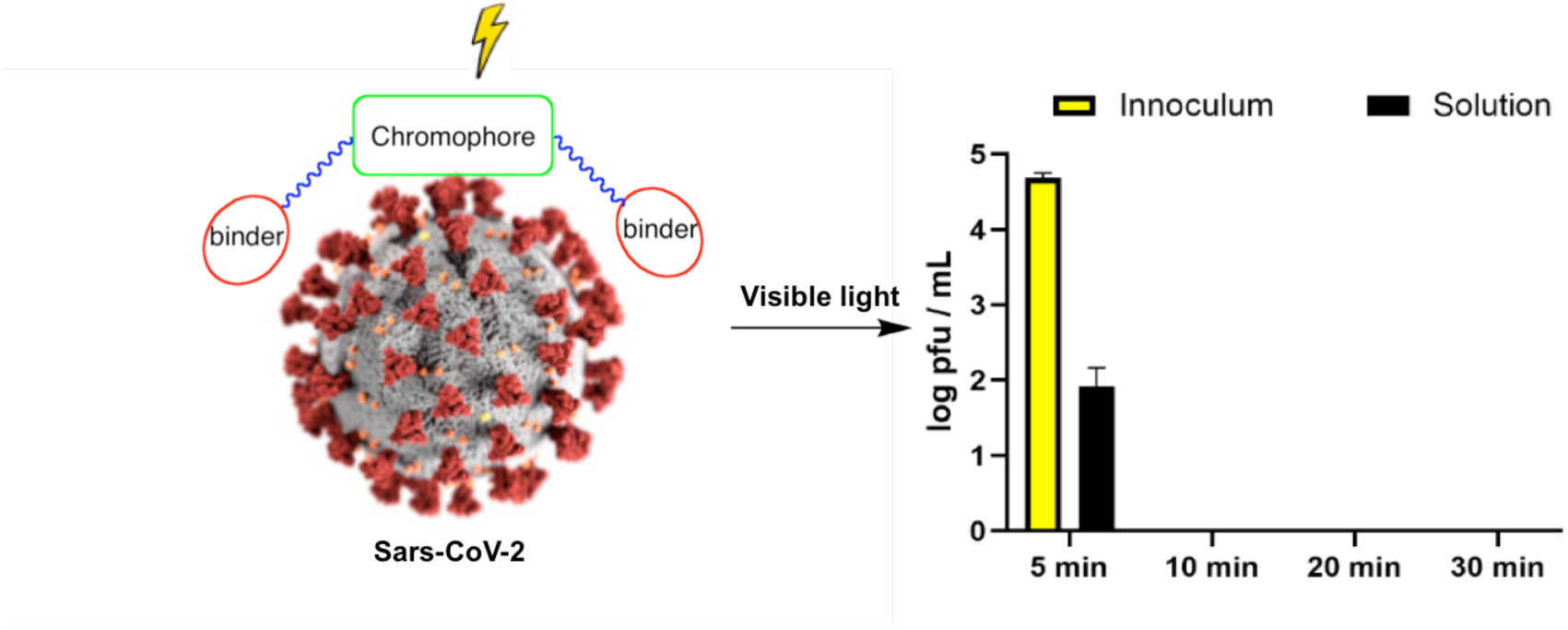

## Footnotes

1 We had to pipette up and down vigorously.

