## Supporting Information for "Rapid and Effective Inactivation of SARS-CoV-2 by a Cationic Conjugated Oligomer with Visible Light: Studies of Antiviral Activity in Solutions and on Supports"

**Supporing Information**

### TABLE OF CONTENTS

Figure S1……….........................................................……………………………..S2

Figure S2…….........................................................……………………………..S2

Experimental Details…………..…….………………………………………..S3-S5

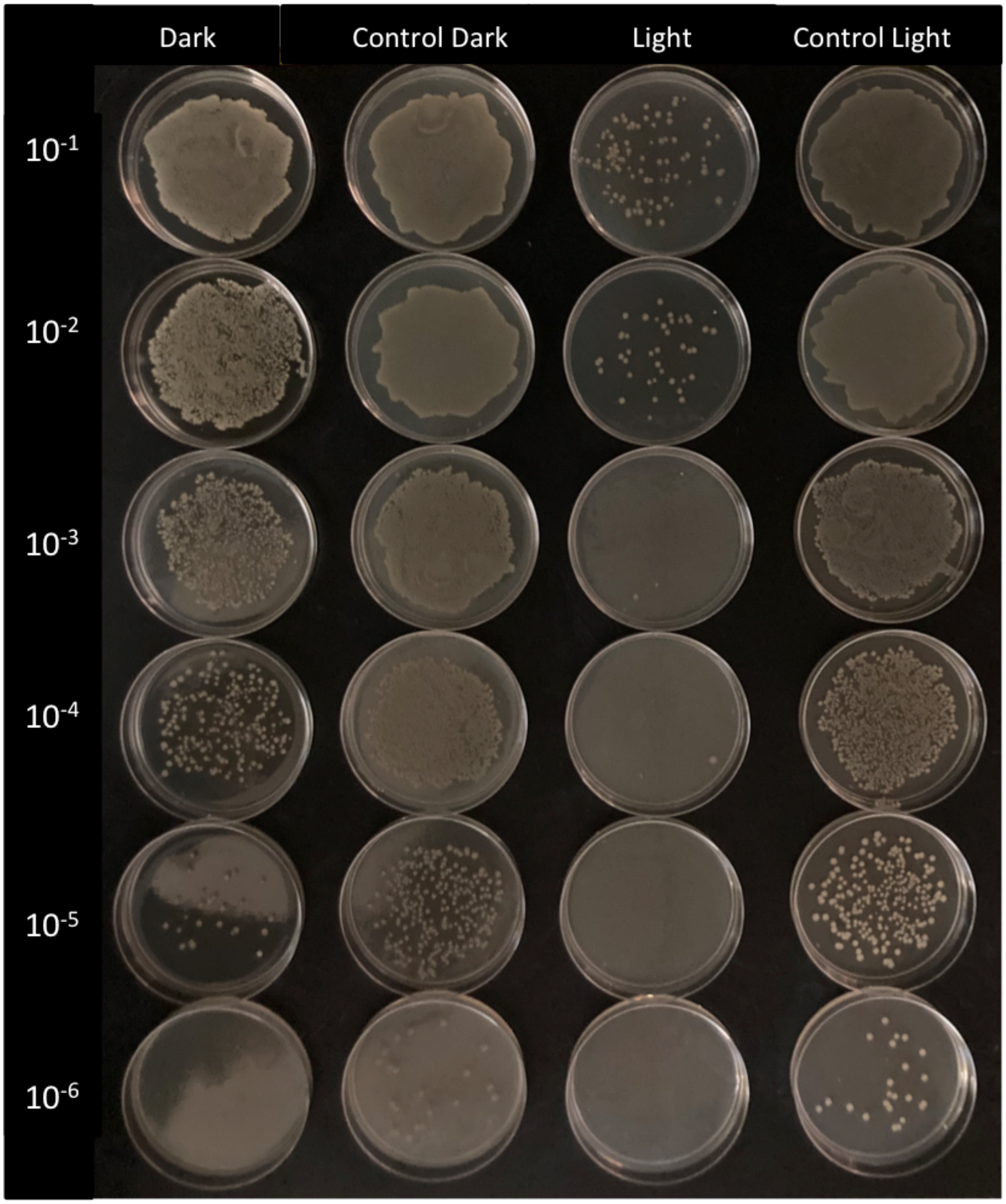


Figure S1. Antimicrobial activity of 1 (10 μg/mL) against E. coli upon exposure to cool white light for 5 min.

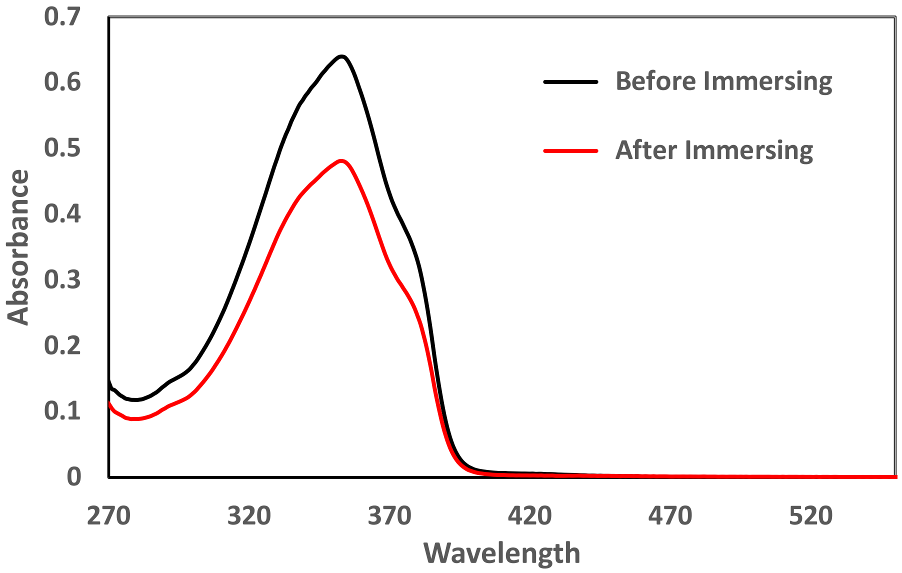


Figure S 2. UV Vis spectra of compound 1 (10 µg/mL) before and after glass fiber immersed.

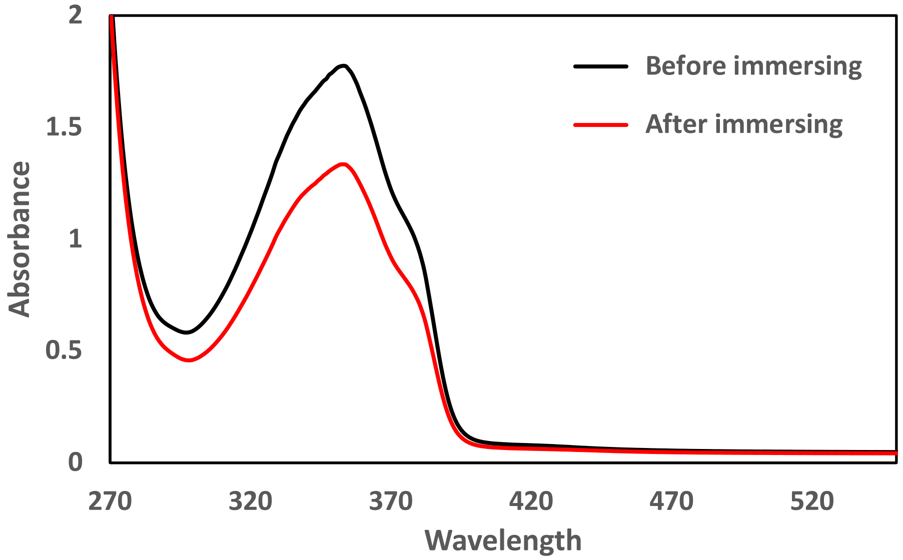


Figure S3. UV Vis spectra of compound 1 (25 µg/mL) before and after commercial wipe paper immersed.

**2,5-bis[ethynyl(trimethylsilyl)] thiophene:**

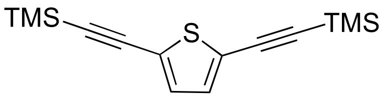


A mixture of 2,5-Dibromothiophene (2.4g, 10.0 mmol), bis(triphenylphosphine)palladium(II) chloride (0.704 g, 1.0 mmol) and copper(I) iodide (0.380 g, 2 mmol) in THF/$Et_3N$ (40mL:20mL) was deoxygenated for 30 min. Trimethylsilyl acetylene (4.92 g, 50.0 mmol) was slowly added and mixture was stirred for 3 days at ambient temperature. The reaction mixture was filtered through a celite pad, and the pad was rinsed with $CH_2Cl_2$.The filtrate was washed with brine and the organic layer was dried with ($Na_2SO_4$), filtered, and concentrated. The residue was purified by column chromatography to afford 2,5-bis[ethynyl(trimethylsilyl)] thiophene. 2.3 gr yield 83%. $^1$H NMR ($CDCl_3$, 300 MHz) δ 0.24 7.04 (s, 2H), (s, 18H); $^{13}$C NMR ($CDCl_3$) δ 132.29, 124.47, 99.90, 96.86, -0.23.

**2,5-diethynylthiophene:**

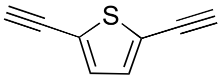


2,5-bis-[(Trimethylsilyl)-ethynyl]-thiophene (1.38 g, 5.0 mmol) was dissolved in methanol (80 mL) and the solution was degassed with nitrogen. To this solution, 0.4 mL of a 0.5 M KOH solution was added, and the mixture was stirred at room temperature for 3 hours. After reaction completed, 200 mL water was added and the mixture was extracted with hexane, the organic phase dried on $Na_2SO_4$, filtered, and the solvent was removed under reduced pressure and used immediately for next step. The desired product was obtained 600 mg, yield 91%. $^1$H NMR ($CDCl_3$, 300 MHz) δ 7.13 (s, 2H), 3.36 (s, 2H); $^{13}$C NMR (CDCl3, 75 MHz) δ 132.8, 123.8, 82.3, 76.4.

**1-(3-bromopropoxy)-4-iodobenzene**

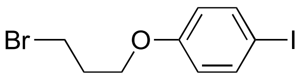


1,3-dibromopropane (24 mL, 363.62 mmol) was added to a solution of 4-iodophenol (4g, 18.18 mmol) and potassium carbonate (10g) in dry acetone (120mL) in a flask and mixture was refluxed at 60 °C under nitrogen for 24 hours. After completion of the reaction, DCM and water was added to the reaction mixture. The organic layer was collected and dried over anhydrous magnesium sulphate. After removal of dichloromethane in vacuo, the residue was purified by silica column chromatography using hexane and ethyl acetate as eluent. 4.4 gr, yield 71%. $^1$H NMR (300MHz, $CDCl_3$), δ: 7.57 (d, $J$ = 9.0 Hz, 2H), 6.70 (d, $J$ = 9.0 Hz, 2H), 4.05 (t, J = 6.0 Hz, 2H), 3.60 (t, $J$ = 6.6 Hz, 2H), 2.31 (m, $J$ = 2H).

**2,5-bis((4-(3-bromopropoxy)phenyl)ethynyl)thiophene**

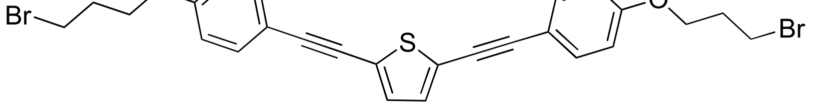


140 mg (0.2 mmol) of $Pd(PPh_3)_2Cl_2$ and 152 mg (0.8 mmol) of CuI were added to a deoxygenated solution of 620 mg (1.82 mmol) of 1-(3-bromopropoxy)-4-iodobenzene (compound **5**) and 529 mg (4 mmol) of

2,5-diethynylthiophene (compound **3**) in 20 mL of $CHCl_3$/ $(iPr)_2NH$ and stirred at room temperature under nitrogen for 3 days. The solvent was removed and the solid was purified by flash chromatography on silicagel with $CHCl_3$ to yield compound. 496 mg, yield 49%. $^1$H NMR (300MHz, $CDCl_3$): 7.46 (d, $J$ = 8.7 Hz, 4H), 7.11 (s, 2H), 6.88 (d, $J$ = 8.7 Hz, 4H), 4.13 (t, $J$ = 5.8 Hz, 4H), 3.61 (t, $J$ = 6.4 Hz, 4H), 2.33 (m, 4H).

**3,3'-(3,3'-(4,4'-(thiophene-2,5-diylbis(ethyne-2,1-diyl))bis(4,1-phenylene))bis(oxy)bis(propane-3,1-diyl))bis(1-methyl-1H-imidazol-3-ium) bromide**

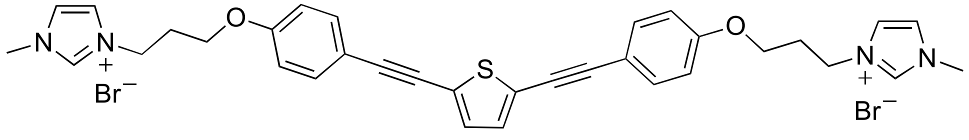


To a suspension of 486 mg (0.87 mmol) compound in 6 mL MeCN, 162 mg (1.97 mmol) N-methyl imidazole in 6 mL MeCN was added. After completion of addition, temperature was increased to 85 °C and refluxed for 3 days. After reaction was completed, solvent was evaporated in vacuo. The reaction mixture was sticky and viscos. To reaction mixture 15 mL $CHCl_3$ was added and evaporated, then, diethyl ether (15 mL) was added and evaporated and rxn mixture became solidified. After solidification, 10 mL DCM was added and stirred for 1 hr. Liquid (DCM) part of the solution decanted and repeated 2 more times. 592 mg, yield 82% $^1$H NMR (300 MHz, DMSO-$d_6$): 9.18 (s, 2H), 7.82 (s, 2H), 7.72 (s, 2H), 7. 52 (d, $J$=8.6 Hz, 4H), 7.34 (s, 2H), 7.96 (d, $J$=8.6 Hz, 4H), 4.35 (t, $J$=6.8 Hz, 4H), 4.07 (t, $J$=5.8 Hz, 4H), 3.84 (s, 6H), 2.29 (t, $J$=6.2 Hz, 4H), $^{13}$C NMR (100 MHz, DMSO-$d_6$): 59.35, 137.23, 133.52, 133.09, 124.08, 122.92, 115.45, 114.01, 94.73, 81.25, 65.28, 46.83, 36.24, 29.40. TOF MS ES+: calculated: 721.08 found 721.0829.

**4,4'-(3,3'-(4,4'-(thiophene-2,5-diylbis(ethyne-2,1-diyl))bis(4,1-phenylene))bis(oxy)bis(propane-3,1-diyl))bis(1-hexyl-1,4-diazoniabicyclo[2.2.2]octane) bromide**

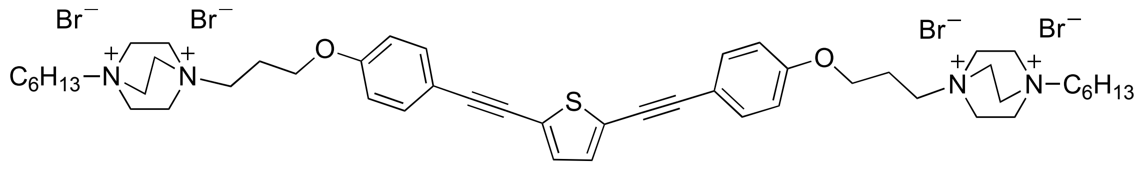


A solution of 253 micromol of DABCO in 1 mL of DMA was added to a solution of 89 micromol of compound **6** in 2 mL of DMA and the mixture solution was stirred at 110 °C for 24 h. The resulting precipitates was collected by filtration and washed with $CHCl_3$.
